# Modeling the seasonal variation of windborne transmission of porcine reproductive and respiratory syndrome virus between swine farms

**DOI:** 10.1101/2023.07.07.547225

**Authors:** Seunghyun Lim, Andres M. Perez, Kaushi S. T. Kanankege

## Abstract

Modeling windborne transmission of aerosolized pathogens is challenging. We adapted an atmospheric dispersion model named Hybrid Single-Particle Lagrangian Integrated Trajectory model (HYSPLIT) to simulate windborne dispersion of porcine reproductive and respiratory syndrome virus (PRRSv) between swine farms and incorporated the findings into an outbreak investigation. The risk was estimated semi-quantitatively based on the cumulative daily deposition of windborne particles, and the distance to closest emitting farm with an ongoing outbreak. Five years of data (2014 : 2018) were used to study seasonal differences of deposition thresholds of the airborne particles containing PRRSv and to evaluate model in relation to risk prediction and barn air filtration. When considered the 14-day cumulative deposition, in Winter, above threshold particle depositions would reach up to 30 km from emitting farms with 84% of them being within 10km. Long-distance pathogen transmission was highest in Winter and Fall, lower in Spring, and least in Summer. The model successfully replicated the observed seasonality of PRRSv where Fall and Winter posing a higher risk for outbreaks. Reaching the humidity and temperature thresholds tolerated by the virus in Spring and Summer reduced the survival and infectivity of aerosols beyond 10 -20 km. Within in the data limitations of voluntary participation, when assumed wind as the sole route of PRRSv transmission, the predictive performance of the model was fair with >0.64 AUC. Barn air filtration was associated with fewer outbreaks, particularly when exposed to high levels of viral particles. The study confirms the usefulness of HYSPLIT models as a tool when determining seasonal effects, distances, and inform near real-time risk of windborne PRRSv transmission that can be useful in future outbreak investigations and implementing timely control measures.

## 1. Background

Porcine reproductive and respiratory syndrome virus (PRRSv) is recognized as the costliest endemic swine pathogen affecting the U.S. with an estimated cost of more than $664 million per year due to production losses, treatment costs, and excessive mortality [1,2]. Transmission of PRRSV between farms is primarily from animal movement [3], contact with infected gilts and sows [4,5], and airborne/windborne transmission of aerosolized particles [6, 7]. Two recent reviews by Anderson et al., 2017 [8] and Arruda et. al., 2019 [9] identified many knowledge and gaps related to the aerosol transmission of PRRSv including the challenge in assessing the windborne local area transmission of aerosolized particles containing viable PRRSv in a near real-time manner, and allotting the level of risk.

Virus containing aerosols in animal disease settings could result from sneezing or coughing of infected animals, dried feces, dust, feed, and debris including hair containing the pathogen [10,11]. Windborne transmission occurs when aerosol particles from infected sources generate a plume that can move in both horizontal, vertical and both directions, and reaches to recipient host. Many authors have studied such dispersal in PRRSv and have revealed that windborne transmission could be a possible transmission route for PRRSV. Dee et al. [6] and Otake et al. [7] observed how far PRRSv moved from an infected farm and found viral particles a long distance away from the infected farm (4.7 km and 91.km, respectively). Survival of PRRSv released into the air is largely related to atmospheric conditions and aerosol characteristics. Meteorological and environmental factors including directional winds of low velocity with sporadic gusts, low temperatures, high relative humidity, and low sunlight levels are also suggested to influence the survival of PRRSv in air [4,6,9]. Therefore, when modeling the windborne dispersion of the virus-containing aerosols, considering virological, epidemiological, and meteorological factors is critical.

Atmospheric dispersion models (ADM) such as Hybrid Single-Particle Lagrangian Integrated Trajectory (HYSPLIT) enable simulating the windborne transmission in a near real-time manner [12]. ADMs are computer models that run these ADM algorithms simulate atmospheric dispersion, as well as the chemical and physical processes of aerosolized particles and gases (i.e., plume), to calculate aerosolized particles deposited at various downwind locations. HYSPLIT, which was developed jointly by the Air Research Laboratory of the National Oceanic and Atmospheric Administration (NOAA: https://www.noaa.gov; accessed on 1 January 2019, College Park, MD, USA), and the Australian Bureau of Meteorology. [17-19,20-22]. The HYSPLIT model is available free to registered and non-registered users through the NOAA Air Resource Laboratory (https://www.ready.noaa.gov/HYSPLIT.php) in web, desktop, or LINUX—based formats.

In a previous study, which used a two week period between March and April 2017, we estimated that aerosolized particle emitted from an infected farm could reach susceptible farms within 25 km, with 53.66% being within 10 km [12]. However, the previous case study did not determine the model validity and did not incorporate the tolerance levels of temperature, UV tolerance, and humidity into the simulations and the potential seasonal variations and the protective (or lack thereof) effect of installation of barn air filters were not determined. Moreover, the software user interface used in the Kanankege et. al., 2022 [12] was not widely available compared to the NOAA HYSPLIT modelling software which is available free to registered and non-registered users through the NOAA Air Resource Laboratory (https://www.ready.noaa.gov/HYSPLIT.php) in web, desktop, and LINUX—based formats. Therefore, in this study, we used 4-years of epidemiological data (2014 – 2018) available from the Morrison Swine Health Monitoring Project (MSHMP) of the University of Minnesota [13, 14; https://vetmed.umn.edu/centers-programs/swine-program/outreach-leman-mshmp/mshmp; accessed on 20 June 2021], to address the above gaps and further investigate the deposition thresholds of the airborne particles containing PRRSv. As done for the previous study, we hypothesize that the local windborne spread of PRRSv is semi-quantifiable in relation to the predisposing meteorological and seasonal factors. Therefore, comparing the deposition of windborne aerosolized particles containing PRRSv across the four seasons, and test whether partial or full air filtration could protect the farms against windborne transmission using this semi-quantitative estimation was a secondary goal. We propose that the study presented here contributes to further confirm the use of ADMs in modeling and investigating between farm windborne transmissions of PRRSv as well as other comparable respiratory pathogenic viruses.

## 2. Data and Methods

### 2.1. ADM Modelling Platform

The long-distance, between-farm windborne dispersion PRRSv was modeled using the Hybrid Single-Particle Lagrangian Integrated Trajectory (HYSPLIT) model, which has been previously used to model the trajectories and dispersion of air pollutants [17-22]. The source code of HYSPLIT can be compiled onto a variety of operating systems and computing environments, including Microsoft Windows, Apple OSX, and Linux. The graphical user interface (GUI) is available on the Windows and OSX that provides for less computing experienced users to use HYSPLT easily. However, using a built-in GUI for large datasets is not a good idea because uploading or downloading large numbers of files over the web is slow, and if it fails before completion, the entire upload or download process must be restarted. The Linux executable, on the other hand, enabled us to analyze hundreds of data sets at once in a reasonable amount of time. For example, HYSPLIT is a spatial explicit model that requires an input file included a geographic location, simulation time and date. With the graphical based tool, only one file is created and accepted to execute, but the Linux executable is enabled to process multiple input files by iterating in parallel. Based on this advantage, we used HYSPLIT in the Linux platform for this study.

### 2.2. HYSPLIT-LINUX model inputs

To execute HYSPLT models in LINUX, three input files called CONTROL (control file), SETUP.CFG (setup file) and metrological data are required. The name change of input files are not allowed to and must be in the same directory to run properly. The control file is divided into five sections: initial setup, pollutants parameters, grid setup, sampling setup, and particle parameters. An example of the three files are included as supplement files. The emission simulations were set to start from 5 a.m., everyday, for a period of 24 hours, and the depositions were measured daily (i.e. every 24h). The parameter relevant to PRRSv including epidemiological features of the virus and the survival of the aerosolized virus in relation to key meteorological features were incorporated according to Kanankege et al., 2022 [12]. Specifically, these included aerosol particle diameter, density, release height and quantity, estimates for the maximum time the virus could remain infective in the air, and the estimated decay. In the absence of PRRSv specific values, such as the virus decay in wind, relevant values that were used in existing and validated wind models for Foot and Mouth Disease virus was used [29-30]. HYSPLIT provide additional parameters that can modify to explicitly model the windborne spread of Foot-and-mouth disease virus (FMDV) [26]. The meteorological parameters for FMDV included air temperature (default value: 24 Celsius), relative humidity (default value: 60 %), and viral decay constant (120 mins). The relative humidity was adapted to 50 % because it was suggested by previous studies [27-28]. Air temperature threshold was set to 30 Celsius based on earlier study [27] and concentration of virus particle decrease when the temperature less than 30 Celsius. Finally, the viral half-life parameter was disabled, but the maximum time in air was set to 72 hours from the source term parameter. A dispersal window of two weeks, during which the incursion event i.e., as a result of receiving a sufficient amount of aerosolized particles containing the virus and therefore an outbreak in a susceptible farm was most likely to have occurred, was assumed. Further details on the data collection, choice of parameter values and the choice of two-week period is justified in the previous study [12]. Forward dispersion model runs were performed using the TAPPAS Web API with direct access to HYSPLIT running on a high-performance cloud computing server. The objective of the forward runs was to assign the risk of PRRSv introduction based on the deposition of particles on all farms. The metric used to assess the potential amount of virus being deposited on susceptible farms was the cumulative 14-day deposition.

The maximum altitude was set to 10,000 m above the ground level (m-AGL), i.e., once the top of the model is defined, the aerosolized particles that reach the top are reflected, i.e., bounced back into the model during the simulations. Both ‘dry’ (gravitational) and ‘wet’ (rainfall) deposition were permitted. These deposition parameters include velocity, average weight, A-Ratio, D-Ratio, effective Henry’s constant, in-cloud, and below-cloud [31-32]. In the model runs, the vertical velocity of the particles was defined by meteorological data. For the simplicity of the analysis, the emitting farms were considered to excrete the same amount of the virus every hour throughout the model run time. The particles were transported with the mean wind plus a random component of motion to account for atmospheric turbulence making the cluster of particles expand in time and space. All other settings such as horizontal and vertical mixing coefficients were used at the default settings of HYSPLIT. Further details on Lagrangian models [33], dry deposition of airborne viruses [34], and details on the dry and wet deposition of atmospheric gases and modeling dry and wet deposition on HYSPLIT are found elsewhere [18,22]. The limitations of HYSPLIT and similar ADMs are discussed elsewhere [15-16].

To utilize this tool, users need to be familiar with LINUX skills and the coordinates of the farms they wish to analyze. By following the methods and control setup outlined in our study, users can easily connect to the tool and input the necessary file combinations and latitude-longitude data. The supplement provides further details on the tool’s setup and usage, facilitating the collective analysis of farm data. By incorporating this tool into outbreak investigations, stakeholders can gain timely information on PRRSv transmission dynamics, aiding in targeted interventions and control measures.

### 2.3. Disease data and the partitioning of the study area

As done in the previous study [12], farms with an ongoing PRRSv outbreak were considered as emitting farms and the farm location and filtration data were acquired from MSHMP (database https://vetmed.umn.edu/centers-programs/swine-program/outreach-leman-mshmp/mshmp; accessed on 20 June 2021; [12]). For this study, data was collected from 167 swine farms in Minnesota, which comprised of commercial (54%), far-row-to-wean (20%), farrow-to-finish (15%), multiplier (6%), boar stud (1%), sow (2%), and other (2%) types (see Figure 1). These farms had pig populations ranging from 30 to 6000, with an average of 1975 pigs per farm. The study period was selected to be from 15^th^January 2014 to 26^th^December 2018. We define an outbreak as farms that have been diagnosed as positive for PRRSv and are experiencing and reporting an ongoing status of the disease on the premises in the relevant week, regardless of the PRRSv lineage. Given that the farms may enter, exit, or not submit data for certain weeks, the total number of farms in the monitoring for each week is expected to vary.

**Figure 1.**
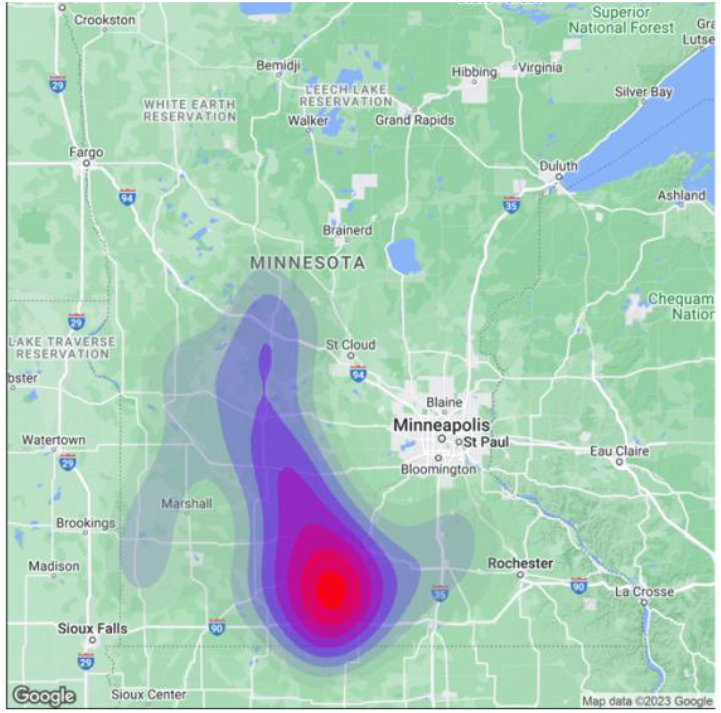
Distribution of the swine 167 swine farms studied; depicted using a kernel smoothing map (Kernel smoothing with coefficient, i.e.bandwidth of 0.05 was estimated using stat_density2d function in R of the Package ‘MASS’ [43]

Having depositions over two weeks from outbreak farms was considered a sufficient period to result in an infection at a down-wind farm, as done in our previous study [12]. There were total of 259 outbreaks, i.e. cases of emitting farms, in the 5-year study period: between 15^th^ January 2014 – 26^th^ December 2018. This resulted in 3766 simulations of particle emission and deposition (i.e., 14 days x 259 = 3626) (see Table S1 in the supplemental material). In the study area, at a given two-week period, there were between 16 – 56 emitting farms.

### 2.4. Wind data

Metrological data is one of the required components for using HYSPLIT to simulate and analyze the transport and dispersion of air pollutants. The HYSPLIT program can be run with a variety of metrological data, including reanalysis data from the National Centers for Environmental Prediction (NCEP) which was used for this study (ftp://arlftp.arlhq.noaa.gov/pub/archives/reanalysis, <u>accessed on 10 August 2022</u>). When using NCEP reanalysis data, HYSPLIT calculates the movement of pollutants over time using information from the reanalysis dataset on atmospheric winds and temperatures. This enables a realistic simulation of pollutant transport and dispersion while accounting for the effects of weather patterns and variability.

### 2.5. HYSPLIT-LINUX model outputs

The key outputs of the models are quantitative maps of the dispersed particle on the terrain over a two-weeks period, (plume deposition per square area mass/m^2^), i.e., the ‘footprint’ of aerosol deposition. The usual HYSPLIT deposition output results in the mass of gases and particles (i.e., plume) per square meter. The daily and cumulative deposition values over the 14-day period were regarded as alternative approximate risk values indicating the likelihood of the introduction of the virus into susceptible farms. The mapped maximum particle deposited at each of the susceptible farm locations was extracted by intersecting the farm locations with the output maps, using s2_closest_feature from the s2 package [38]. The particles deposited at each susceptible farm during the two-week period were used to represent the “exposure hazard” (i.e., potential for pathogen introduction via aerosols). In this retrospective modeling exercise, it is challenging to determine an infectious dose that is sufficient to cause the windborne disease.

### 2.6. Data analysis: seasonality and barn air filtration

To determine a threshold deposition for infection, we presumed that farm-to-farm aerosol infection is uncommon and that only the higher deposition values might result in a farm becoming infected by this route. To estimate this, we produced histograms of the deposition values for the three metrics and applied natural breaks/Jenks classification (Jenks, 1967) for the cumulative, median, and maximum daily deposition. The frequency distributions were divided into three classes using two natural breaks (i.e., Jenks). We presumed that a very low exposure dose is unlikely to result in infection, and to convert the exposure hazard and may have the potential risk of contracting the infection, we applied an infecting dose threshold [35]. As discussed in Kanankege et al., 2022 [12] study on HYSPLIT applicable parameters for PRRSv, in the absence of field data relevant to the farms in the study, we determined these thresholds by examining the histograms of the cumulative, median daily, and maxi-mum daily concentrations. Applying these thresholds to the susceptible farms, we thus were able to classify them as receiving a potentially sufficient infecting dose or not; the same farm that was classified as a ‘Farm at risk’. The distance to farms at risk from the closest emitting farm that is upwind were calculated. In this distance calculation, a farm may be categorized as at risk under the threshold for cumulative deposition over the 14-day period. To compare the deposition of airborne particles containing PRRSv across the seasons, the cumulative 14-day deposition on a farm was summarized by four seasons. The seasons were defined as Spring (3/21 through 6/20), Summer (6/21 through 9/20), Fall (9/21 through 12/20), and Winter (12/21 through 3/20) [36]. After 14-days of deposition, applying natural break thresholds to the three-deposition metric of 14-day cumulative, median, and maximum daily deposition, the following values were determined as deposition thresholds. Natural break of the distribution of the depositions were used to assign the threshold to determine high-risk farms, i.e. farms were classified as high risk farms if their exposure to the particles exceeded the first threshold value, indicating a sufficient number of particles to pose a risk for airborne introduction of the pathogen.

To determine whether air filtration systems could effectively reduce the risk of infection among pigs and prevent the spread of the virus, we incorporated filtration status of the farm when further analyzing particle deposition and new cases While the filtration status of some farms changed over time, the analysis included filtered (n=49), partial (n=15), not filtered (n=56), and unknown (n=47) farms. Farms with unknown filtration status were excluded from this comparison, and the transmission was considered primarily due to windborne transmission.

### 2.6. Model cross-validation: AUC, Sensitivity, and Specificity

The receiver operating characteristic (ROC) curve and the area under the curve (AUC) are widely used in various research domains to evaluate the performance of predictive models. The ROC curve is a graphical representation of the relationship between the sensitivity (true positive rate) and specificity (true negative rate) of a model at different decision thresholds. The AUC is a summary statistic that quantifies the overall performance of the model across all possible decision thresholds, ranging from 0.5 (chance performance) to 1.0 (perfect discrimination). To evaluate the performance of predictive models, the result of the 14-day cumulative concentration of farms with and without a new outbreak was used. Out of 269 total instances, those that have at least one newly infected farm were chosen, resulting in 118 cases for analysis. The 14-day plume concentration value for only uninfected farms was extracted and identified new outbreak status for each farm. Then all the 14-day cumulative concentration data were grouped by season and compared the farms with a new outbreak to those without using an incremental threshold for each of the four seasons.

## 3. Results

### 3.1. HYSPLIT-LINUX Deposition Thresholds and Seasonality

Deposition thresholds determined at the 1st natural break of the particle deposition varied by season (Figure 2 and Table 3). Across all three indices of 14-day cumulative deposition, median daily deposition, and maximum daily deposition; both spring and summer had higher deposition thresholds compared to fall and winter. Yet the particle travel distances with wind were higher in both Spring and Summer (Table 4). Whereas, in the Fall and Winter, the deposition thresholds were low and particle travel distances were less as well (Table 4). This resulted in highest numbers of above threshold cumulative and median particle deposition at distances away from an emitting farm to occur in Winter months (i.e. December – March), making Winter the highest risk period for long distance windborne transmission of the PRRSv (Tables 4.1 and 4.2). This followed by Fall, Spring, and Summer at the least risk for long distance transmissions via wind.

**Figure 2.**
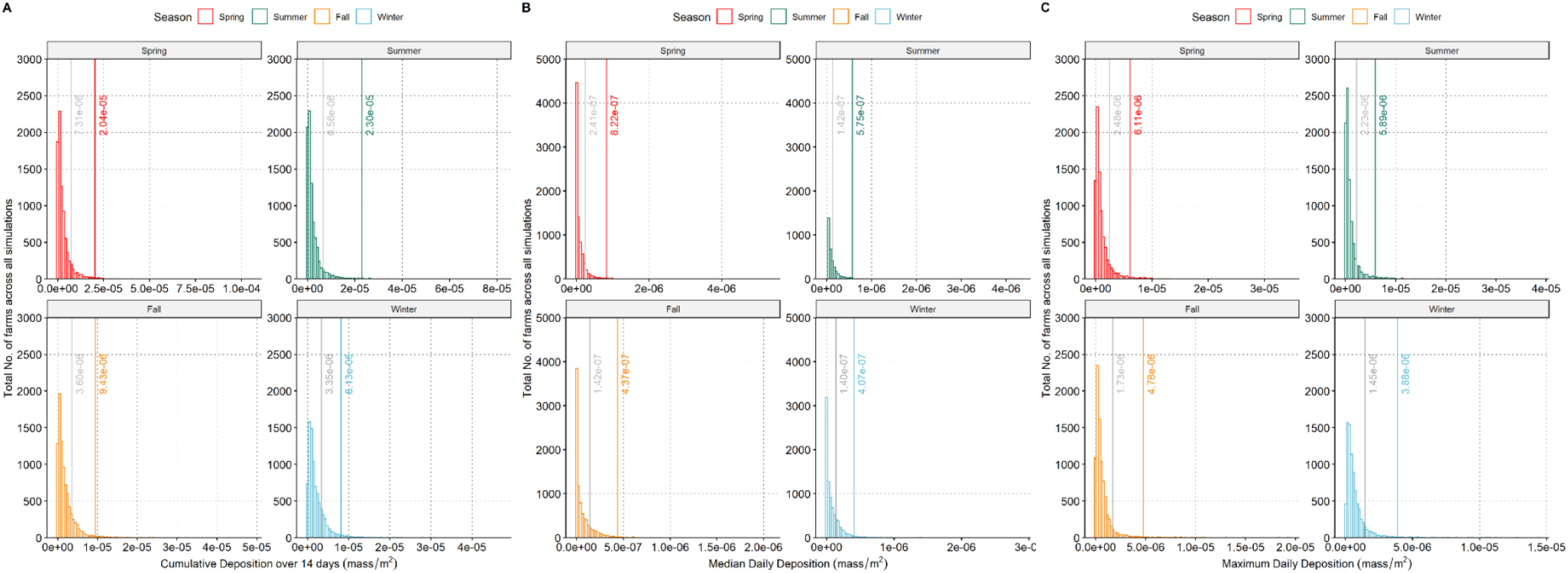
Panel A) Fourteen-day cumulative deposition of windborne particles, Panel B) Median daily deposition, and Panel C) Maximum daily deposition on swine farms by season; over the 5-year period (y-axis: total number of farms across all simulations, x-axis: mass/m^2^). The seasonal thresholds set at natural breaks 1 and 2 of the data distrbution is indicated in each figure.

**Table 3.**
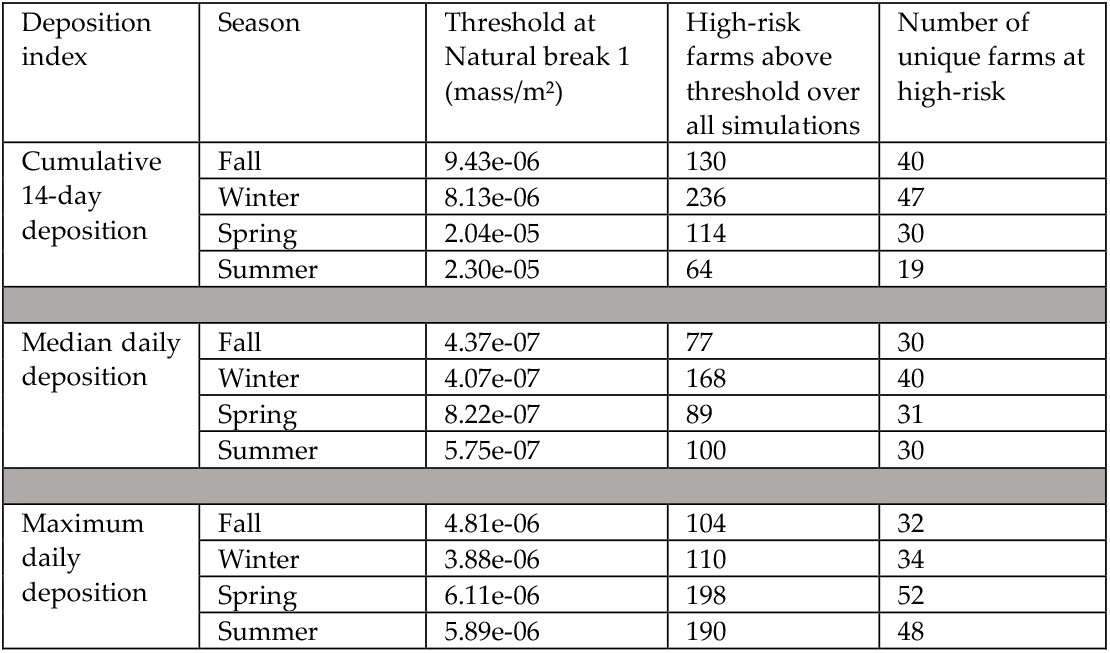
Particle deposition thresholds for three indices: 14-day cumulative, median daily, and maximum (mass/m2), and the number of farms ≥ the threshold at each season for all simulations over the 5-year study period.

**Table 4.**
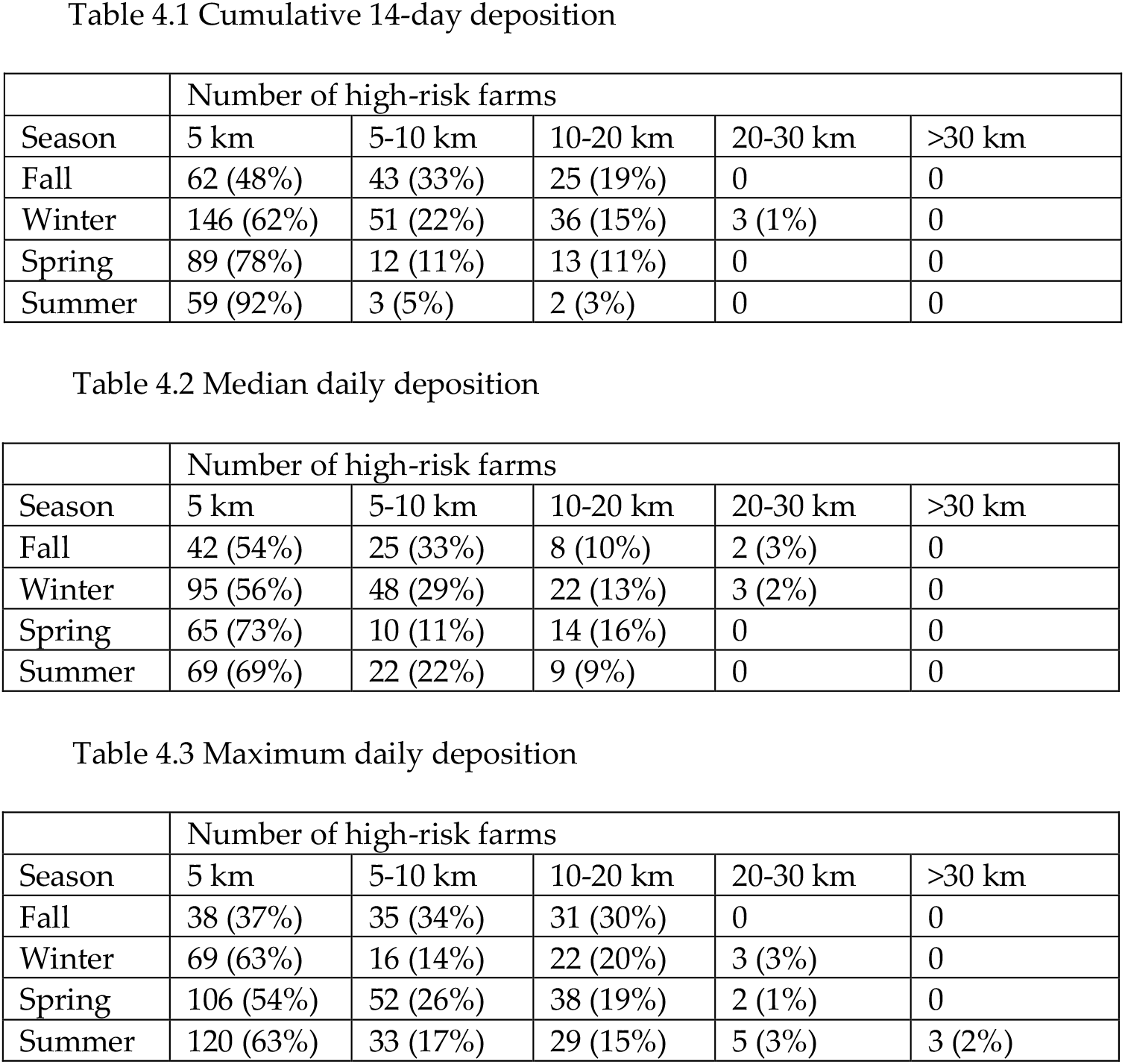
Distance to high-risk farms from an emitting farm by season.

When the 14-day cumulative deposition data, during the Winter season, particles were found to reach distances of up to 30 km, with 62% of them being within 5 km. In the Fall season, particles reached up to 20 km, with 48% within 5 km. For the Spring season, particles reached up to 20 km, with 78% within 5 km. Similarly, during the summer season, particles reached up to 20 km, with 92% within 5 km. While occasional long-distance depositions beyond 20 km were observed in the summer season according to the maximum daily index (Table 4.3), the cumulative deposition analysis indicated that only 8% of particles persisted beyond 5 km. This is because simulations would cease the particles existence when the temperature or humidity thresholds of the PRRSv were reached. In contrast, the Fall and Winter seasons exhibited cumulative deposition above the threshold at distances of 10 km and beyond. The Fall season had the highest percentage (19%), followed by Winter (16%). Consequently, these observations indicate that Fall and Winter seasons may pose a higher risk for long-distance pathogen transmission compared to Spring and Summer seasons.

### 3.4. The effect of barn air filtration

The analysis showed that the probability of having a PRRSv outbreak was similar between filtered and non-filtered farms when the farm was not exposed to a critical level of the virus in the air (as defined by the threshold) (61.5% vs. 63%, Figure 3). This could be due to various factors such as virus transmission through movements or re-emergence of the virus in an already infected farm. However, when the farm was exposed to a critical level of PRRSv in the air, the probability of having an outbreak was almost four times higher for non-filtered farms compared to filtered farms (3.7/0.96 = 3.85 Figure 3). The findings suggest that farms that implemented filtration measures had a lower incidence of reported PRRSv outbreaks compared to farms that were only partially or not filtered, particularly when exposed to high levels of viral particles, and were therefore considered to be at high risk.

**Figure 3.**
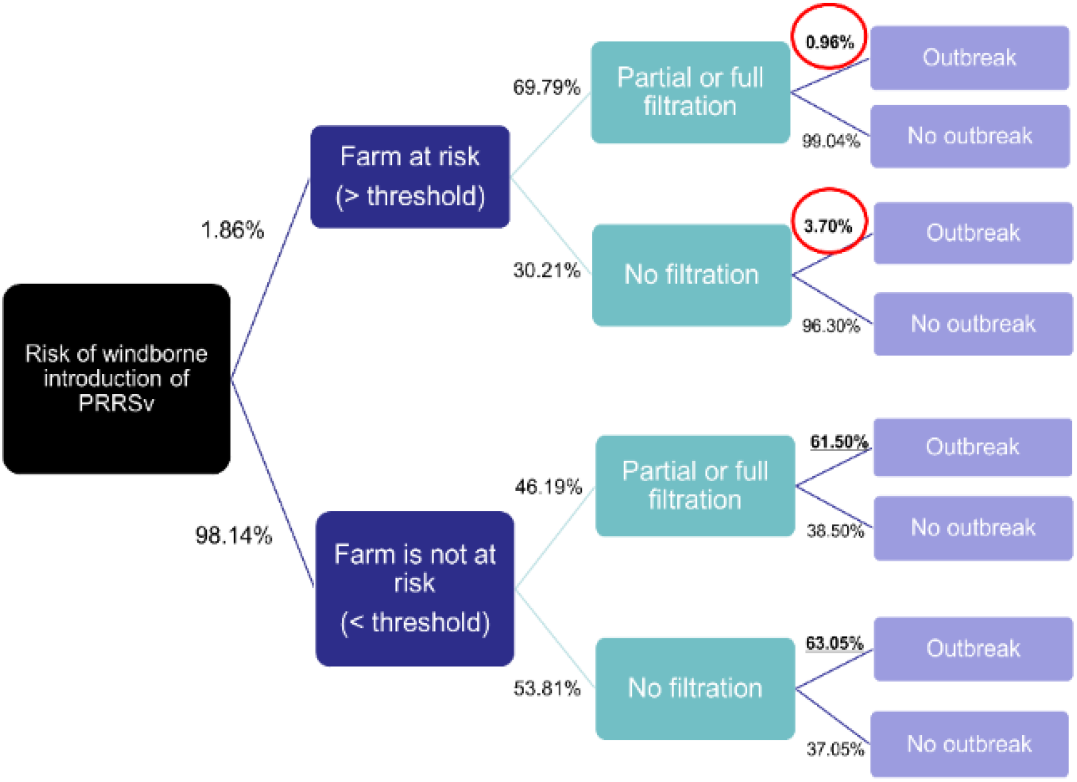
Decision tree analysis comparing the farms at risk, outbreaks, and barn filtration. Farms receiving depositions above threshold were ‘at risk’. Figure adapted from [37]

### 3.4. Model cross-validation: AUC, Sensitivity, and Specificity

To further evaluate the predictive performance of our model, we calculated the AUC value for each season. Our results showed that our model had the highest predictive accuracy during the Winter season and the lowest during the Summer season. The Spring and Fall seasons showed intermediate predictive accuracy. Table 5 summarizes the true positive, false negative, false positive, true negative, sensitivity, and specificity at the two natural break thresholds for each season. The threshold for each season with the highest possible combination value of sensitivity and specificity was also shown in table 5.

**Table 5.**
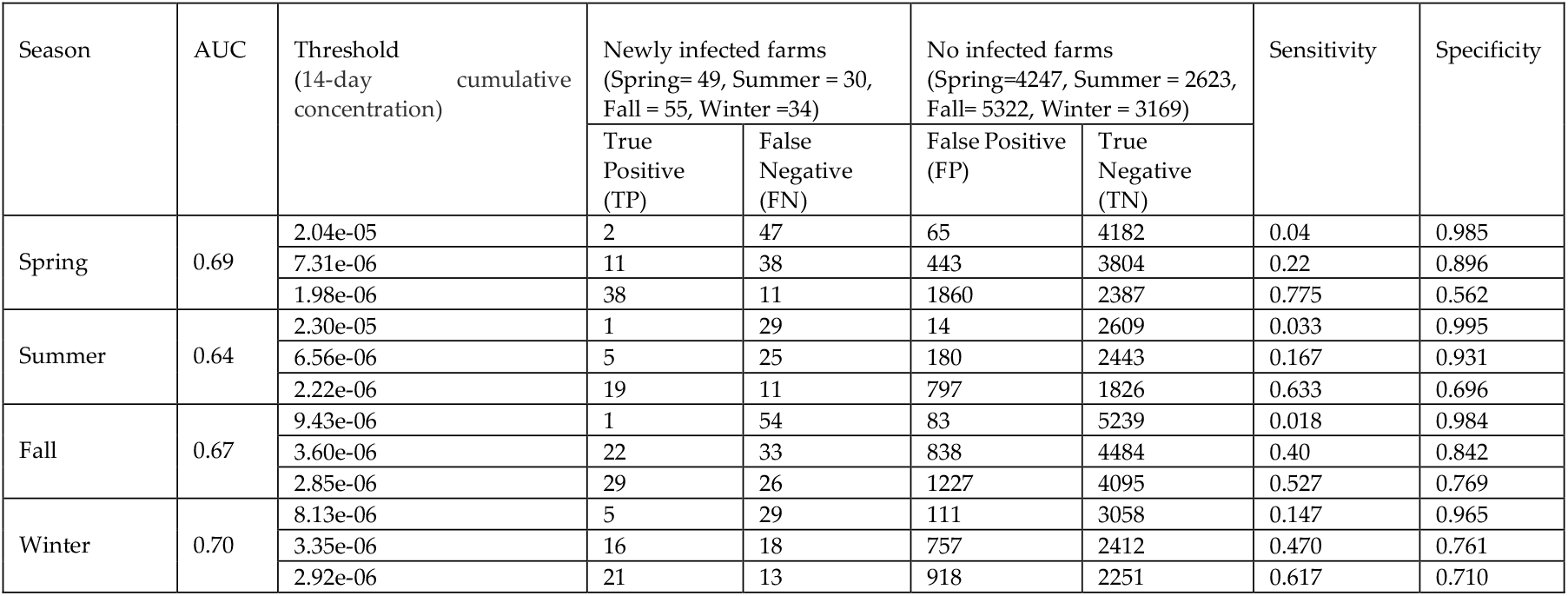
Summary of model performance in predicting the risk of PRRSv outbreak at farms by season.

| Season | AUC | Threshold<br>(14-day cumulative<br>concentration) | Newly infected farms<br>(Spring= 49, Summer = 30,<br>Fall = 55, Winter =34) |  | No infected farms<br>(Spring=4247, Summer = 2623,<br>Fall= 5322, Winter = 3169) |  | Sensitivity | Specificity |
| --- | --- | --- | --- | --- | --- | --- | --- | --- |
|  |  |  | True<br>Positive<br>(TP) | False<br>Negative<br>(FN) | False Positive<br>(FP) | True<br>Negative<br>(TN) |  |  |
| Spring | 0.69 | 2.04e-05 | 2 | 47 | 65 | 4182 | 0.04 | 0.985 |
|  |  | 7.31e-06 | 11 | 38 | 443 | 3804 | 0.22 | 0.896 |
|  |  | 1.98e-06 | 38 | 11 | 1860 | 2387 | 0.775 | 0.562 |
| Summer | 0.64 | 2.30e-05 | 1 | 29 | 14 | 2609 | 0.033 | 0.995 |
|  |  | 6.56e-06 | 5 | 25 | 180 | 2443 | 0.167 | 0.931 |
|  |  | 2.22e-06 | 19 | 11 | 797 | 1826 | 0.633 | 0.696 |
| Fall | 0.67 | 9.43e-06 | 1 | 54 | 83 | 5239 | 0.018 | 0.984 |
|  |  | 3.60e-06 | 22 | 33 | 838 | 4484 | 0.40 | 0.842 |
|  |  | 2.85e-06 | 29 | 26 | 1227 | 4095 | 0.527 | 0.769 |
| Winter | 0.70 | 8.13e-06 | 5 | 29 | 111 | 3058 | 0.147 | 0.965 |
|  |  | 3.35e-06 | 16 | 18 | 757 | 2412 | 0.470 | 0.761 |
|  |  | 2.92e-06 | 21 | 13 | 918 | 2251 | 0.617 | 0.710 |

## 5. Discussion

In this semi-quantitative study, we adapted NOAA-HYSPLIT-LINUX to model windborne dispersion of aerosolized PRRSv between farms and determined the varying levels of windborne transmission of PRRSv containing particles and the influence of barn air filtration to potentially prevent the disease. In a previous study, which used a two-week example three-week period between March and April 2017, we estimated that aerosolized particle emitted from an infected farm could reach susceptible farms within 25 km, with 53.66% of the farms being within 10 km. Extending the objectives further, in this 5-year study where we incorporated virus survival and decay characteristics into the wind models. With this modeling exercise we confirm the usefulness of HYSPLIT models as a tool when determining seasonal effects, distances, and inform near real-time risk of windborne PRRSv transmission that can be useful in future outbreak investigations and planning control measures such as renewal of air filtration or determining the ineffective ones.

Moreover, this tool holds significant potential in determining the risk differences among seasons. Previous studies have consistently demonstrated seasonal variations in PRRSv incidence [39,40]. Typically, PRRSv in the Midwest of the USA exhibits higher incidence rates during fall and winter, with reduced activity in spring and summer. The simulation of airborne transmission in Table 4 provides evidence to support the observation of seasonality in PRRSv outbreaks. Results indicated that the distance from an emitting farm to a susceptible farm that could receive above threshold ‘infectious’ number of particles could vastly vary by season. Specifically, when considered the 14-day cumulative deposition, in summer particles would reach up to 20 km with 92% being with in 5km, in Fall up to 20 km with 48% within 5km, in Winter up to 30 km with 62% within 5km, and in Spring up to 20 km with 78% within 5km. The temperature and humidity during summers influenced the viral particles to not survive alive and infective long enough beyond 10 – 20 km and therefore the risk of pathogen introduction was determined to be low compared to Fall and Winter seasons (Table 4). The study also suggested that farms that were at high-risk based on above threshold particle depositions, but were filtered, tend to have less reported outbreaks compared to partially or non-filtered farms.

Barn air filtration was associated with fewer outbreaks, particularly when exposed to high levels of viral particles. This provides valuable insights into the effectiveness of filtration systems in swine farming areas. Modeling the risk of windborne transmission using an ADM like HYSPLIT offers several advantages over short distance experiments conducted in the field. an experimental study s reported that air-filtration systems were effective in reducing aerosol transmission of PRRSv among pigs housed in chambers connected by an airduct [42]. Another 4-year field study reported that farms with air filters compared to non-filtered controls were less likely to end up with PRRSv, supporting further application in the field especially in pig dense areas [44]. While field experiments provide valuable data on localized transmission dynamics, they may not fully capture the long-range airborne transmission patterns that occur between farms located at a significant distance from each other. By integrating data from ADM modeling and a comprehensive dataset of multiple participant farms from a larger area, we can better assess the cumulative risks associated with long-range transmission, particle deposition as a function of several outbreak farms down wind and evaluate the efficacy of air filtration systems in mitigating these risks. Within in the data limitations of voluntary participation, when assumed wind as the sole route of PRRSv transmission, the predictive performance of the model was fair with Area under the ROC curve (AUC) ranging between 0.64 – 0.70. The average specificity was 0.84 across the seasons indicating the modeling tool could support “ruling out” the potential of windborne introduction of the pathogen, if used in an outbreak investigation, when the farm status of the surrounding farms were available for the simulation. The average sensitivity was 0.34, however, this is also relatable to the fact that cross-validation was done for the specific weeks where there were newly infected farms, which were limited.

However, it is crucial to understand that atmospheric dispersion modeling alone cannot conclusively confirm aerosol transmission as the cause of a new outbreak on a farm, hence the interpretation of should be done within the context of the study. Demonstrating through dispersion modeling that a high concentration of the virus could have been deposited only suggests windborne transmission as a possible pathway for pathogen introduction. Dispersion modeling is valuable in ruling out aerosol dispersion between farms as a transmission route when the results indicate deposition is not feasible or at a low concentration [32]. By solely considering airborne transmission, our study might have underestimated the complete risk and dynamics of PRRSv transmission in the population. Future studies should strive to incorporate multiple transmission routes for a more comprehensive understanding of PRRSv transmission dynamics. While we incorporated important environmental factors such as temperature, relative humidity and decay rate, as they are known to impact the stability and survival of PRRSv particles in the atmosphere. However, we did not account for other influential factors such as UV radiation, which can also influence the viability of virus particles [37,41,42]. There is limited data available on the PRRSv sensitivity to UV radiation yet neglecting these additional factors may limit the comprehensive understanding of PRRSv transmission dynamics and could potentially lead to underestimation or overestimation of transmission risks. Another limitation is the difficulty in estimating farm-to-farm transmission in an endemic setting, especially with a voluntary testing program. Due to the voluntary nature of the program, it is impossible to determine the PRRSv status of all farms. PRRSv transmission dynamics involve viral fadeout and reintroduction, influenced by factors such as larger herds in pig-dense regions and continuous introduction of infectious stock. To enhance model prediction accuracy and risk estimation, having data from all pig farms in a given area would be valuable. Also, modeling the windborne between farm transmission of PRRSv in Midwestern and Southeastern regions of the U.S. where swine farming is prominent would shed more light on the effect of different terrain, wind and climatological factors, management, airfiltration cost-effectiveness, on the windborne transmission of PRRSv. Such comprehensive studies would inform more effective prevention and control strategies.

## 6. Conclusions

The HYSPLIT model was adapted to simulate the windborne dispersion of PRRSv and investigated its transmission dynamics between swine farms. The findings provide valuable insights into the seasonal differences and deposition thresholds of airborne particles carrying PRRSv. Winter and Fall seasons were identified as periods with a higher risk of long-distance transmission, while Spring and Summer exhibited lower transmission potential due to reduced survival and infectivity of aerosols. The model demonstrated fair predictive performance, indicating its usefulness in informing near real-time risk assessment of windborne PRRSv transmission and guiding outbreak investigations. Implementing filtration measures showed promise in reducing the incidence of reported PRRSv outbreaks, especially in the presence of high viral particle levels. Overall, this five-year simulation study contributes to our understanding of windborne transmission dynamics and provides valuable insights for implementing effective control measures to mitigate the spread of PRRSv. Future research should focus on refining the model, incorporating additional factors influencing transmission, and improving data collection to further enhance our ability to predict and manage PRRSv outbreaks.

## Supporting information

ASCDATA file

CONTROL file

SETUP file

## Supplementary Material

The following supporting information can be downloaded at: XX

Table S1.

SETUP.cfg file.

ASCDATA.cfg file.

CONTROL file.

## Author Contributions

Conceptualization, methodology, formal analysis, and validation: K.S.T.K., S.L. and A.M.P. Investigation, K.S.T.K., and S.L.; Resources, A.M.P.; Data Curation, S.L. and K.S.T.K., and S.L.; Writing—Original Draft Preparation, K.S.T.K. and S.L.; Writing—Review & Editing, A.M.P. and K.S.T.K.; Visualization, S.L.; Supervision, K.S.T.K. and A.M.P.; Project Administration, K.S.T.K.; Funding Acquisition, A.M.P. and K.S.T.K. All authors have read and agreed to the published version of the manuscript.

## Institutional Review Board Statement

Not applicable.

## Informed Consent Statement

Not applicable.

## Data Availability Statement

The data are not publicly available due to privacy concerns. The data presented in this study are available on request from the Morrison Swine Health Monitoring Project (MSHMP) of the University of Minnesota (restrictions apply).

## Acknowledgments

The authors gratefully acknowledge the NOAA Air Resources Laboratory (ARL) for the provision of the HYSPLIT transport and dispersion model and/or READY website (https://www.ready.noaa.gov, accessed on 20 January 2020) used in this publication.

## Abbreviations

ADM: Atmospheric dispersion models
HYSPLIT: Hybrid Single-Particle Lagrangian Integrated Trajectory models
m-AGL: meters above ground level
NAM: North American Mesoscale Forecast System
PRRSv: porcine reproductive and respiratory syndrome virus
TAPPAS: Tool for Assessing Pest and Pathogen Aerial Spread.

