## Supplementary material for "Modeling the seasonal variation of windborne transmission of porcine reproductive and respiratory syndrome virus between swine farms": ASCDATA file

**ASCDATA.CFG**

The model implementation utilized the sample ASCDATA.CFG file. For a comprehensive understanding of the parameter's definition, please refer to the HYSPLIT user guide provided at the following link: https://www.ready.noaa.gov/hysplitusersguide/S444.htm

```
-90.0  -180.0
1.0  1.0
180  360
2
0.2
'.'
```
