## Supplementary material for "Modeling the seasonal variation of windborne transmission of porcine reproductive and respiratory syndrome virus between swine farms": CONTROL file

The model implementation utilized the sample Control file. For a comprehensive understanding of the parameter's definition, please refer to the HYSPLIT user guide provided at the following link: https://www.ready.noaa.gov/hysplitusersguide/S310.htm

```
0 0 0 0
1
40.0 -90.0 50.0
48
0
10000
1
met_data
met_data_name
1
part
1.0
1.0
0 0 0 0
1
40.0 -90.0
1.0 1.0
180.0 260.0
./
output.bin
1
50
0 0 0 0
1 0 0 0
0 24 0
1
0 0 0
0 0 0 0 0
0 0 0
0
0
/
```
