## Supplementary material for "Modeling the seasonal variation of windborne transmission of porcine reproductive and respiratory syndrome virus between swine farms": SETUP file

**SETUP.CFG file**

The model implementation utilized the sample SETUP.CFG file. For a comprehensive understanding of the parameter's definition, please refer to the HYSPLIT user guide provided at the following link: https://www.ready.noaa.gov/hysplitusersguide/S410.htm

```
&SETUP
tratio=0.75,
delt=0.0,
initd=0,
kpuff=0,
khmax=9999,
khinp=0,
numpar=2500,
maxpar=10000,
nbptyp=1,
qcycle=0.0,
efile='',
k10m=1,
kdef=0,
krand=2,
kzmix=0,
kbls=1,
kblt=0,
isot=-99,
idsp=1,
wvert=.FALSE.,
vscale=200.0,
vscales=5.0,
vscaleu=200.0,
hscale=10800.0,
capemin=-1.0,
tkemin=0.001000000,
uratio=5.873300076,
tvmix=1.00,
tkerd=0.18,
tkern=0.18,
kmix0=150,
kmixd=0,
ninit=1,
ndump=0,
ncycl=0,
pinbc='PARINBC',
pinpf='PARINIT',
poutf='PARDUMP',
messg='MESSAGE',
vdist='VMSDIST',
mgmin=10,
conage=24,
gemage=48,
kmsl=0,
kwet=1,
ichem=0,
cpack=1,
cmass=0,
kspl=1,
krnd=6,
frhmax=3.00,
```

```
splitf=1.00,
frhs=1.00,
frvs=0.01,
frts=0.10,
dxf=1.00,
dyf=1.00,
dzf=0.01,
/
```
