## Supplementary material for "Modeling the seasonal variation of windborne transmission of porcine reproductive and respiratory syndrome virus between swine farms": Table S1

The table represents a total of 259 outbreak instances, referring to cases from emitting farms, observed over a 5-year study period from 15th January 2014 to 26th December 2018.

| Case | Week 1 | Week 2 | Number of Participant Farms | No. Infected and Excreting Farms | No. Non-Infected Susceptible Farm | Week 3 | Newly Infected Excreting Farms on week 3 |
|---|---|---|---|---|---|---|---|
| 1 | 1-Jan-2014 | 8-Jan-2014 | 167 | 28 | 139 | 15-Jan-2014 | 0 |
| 2 | 8-Jan-2014 | 15-Jan-2014 | 167 | 28 | 139 | 22-Jan-2014 | 0 |
| 3 | 15-Jan-2014 | 22-Jan-2014 | 167 | 28 | 139 | 29-Jan-2014 | 0 |
| 4 | 22-Jan-2014 | 29-Jan-2014 | 167 | 28 | 139 | 5-Feb-2014 | 0 |
| 5 | 29-Jan-2014 | 5-Feb-2014 | 167 | 29 | 138 | 12-Feb-2014 | 0 |
| 6 | 5-Feb-2014 | 12-Feb-2014 | 167 | 29 | 138 | 19-Feb-2014 | 0 |
| 7 | 12-Feb-2014 | 19-Feb-2014 | 167 | 29 | 138 | 26-Feb-2014 | 0 |
| 8 | 19-Feb-2014 | 26-Feb-2014 | 167 | 29 | 138 | 5-Mar-2014 | 0 |
| 9 | 26-Feb-2014 | 5-Mar-2014 | 167 | 29 | 138 | 12-Mar-2014 | 0 |
| 10 | 5-Mar-2014 | 12-Mar-2014 | 167 | 29 | 138 | 19-Mar-2014 | 0 |
| 11 | 12-Mar-2014 | 19-Mar-2014 | 167 | 29 | 138 | 26-Mar-2014 | 0 |
| 12 | 19-Mar-2014 | 26-Mar-2014 | 167 | 29 | 138 | 2-Apr-2014 | 0 |
| 13 | 26-Mar-2014 | 2-Apr-2014 | 167 | 33 | 134 | 9-Apr-2014 | 0 |
| 14 | 2-Apr-2014 | 9-Apr-2014 | 167 | 32 | 135 | 16-Apr-2014 | 0 |
| 15 | 9-Apr-2014 | 16-Apr-2014 | 167 | 32 | 135 | 23-Apr-2014 | 0 |
| 16 | 16-Apr-2014 | 23-Apr-2014 | 167 | 31 | 136 | 30-Apr-2014 | 0 |
| 17 | 23-Apr-2014 | 30-Apr-2014 | 167 | 29 | 138 | 7-May-2014 | 0 |
| 18 | 30-Apr-2014 | 7-May-2014 | 167 | 28 | 139 | 14-May-2014 | 0 |
| 19 | 7-May-2014 | 14-May-2014 | 167 | 28 | 138 | 21-May-2014 | 1 |
| 20 | 14-May-2014 | 21-May-2014 | 167 | 28 | 139 | 28-May-2014 | 0 |
| 21 | 21-May-2014 | 28-May-2014 | 167 | 25 | 141 | 4-Jun-2014 | 1 |
| 22 | 28-May-2014 | 4-Jun-2014 | 167 | 26 | 141 | 11-Jun-2014 | 0 |
| 23 | 4-Jun-2014 | 11-Jun-2014 | 167 | 26 | 140 | 18-Jun-2014 | 1 |
| 24 | 11-Jun-2014 | 18-Jun-2014 | 167 | 27 | 139 | 25-Jun-2014 | 1 |
| 25 | 18-Jun-2014 | 25-Jun-2014 | 167 | 28 | 139 | 2-Jul-2014 | 0 |
| 26 | 25-Jun-2014 | 2-Jul-2014 | 167 | 26 | 140 | 9-Jul-2014 | 1 |
| 27 | 2-Jul-2014 | 9-Jul-2014 | 167 | 26 | 140 | 16-Jul-2014 | 1 |
| 28 | 9-Jul-2014 | 16-Jul-2014 | 167 | 26 | 141 | 23-Jul-2014 | 0 |
| 29 | 16-Jul-2014 | 23-Jul-2014 | 167 | 24 | 143 | 30-Jul-2014 | 0 |
| 30 | 23-Jul-2014 | 30-Jul-2014 | 167 | 23 | 143 | 6-Aug-2014 | 1 |
| 31 | 30-Jul-2014 | 6-Aug-2014 | 167 | 24 | 143 | 13-Aug-2014 | 0 |
| 32 | 6-Aug-2014 | 13-Aug-2014 | 167 | 23 | 144 | 20-Aug-2014 | 0 |
| 33 | 13-Aug-2014 | 20-Aug-2014 | 167 | 21 | 146 | 27-Aug-2014 | 0 |
| 34 | 20-Aug-2014 | 27-Aug-2014 | 167 | 20 | 147 | 3-Sep-2014 | 0 |
| 35 | 27-Aug-2014 | 3-Sep-2014 | 167 | 19 | 148 | 10-Sep-2014 | 0 |
| 36 | 3-Sep-2014 | 10-Sep-2014 | 167 | 19 | 148 | 17-Sep-2014 | 0 |
| 37 | 10-Sep-2014 | 17-Sep-2014 | 167 | 19 | 148 | 24-Sep-2014 | 0 |
| 38 | 17-Sep-2014 | 24-Sep-2014 | 167 | 19 | 148 | 1-Oct-2014 | 0 |
| 39 | 24-Sep-2014 | 1-Oct-2014 | 167 | 19 | 148 | 8-Oct-2014 | 0 |
| 40 | 1-Oct-2014 | 8-Oct-2014 | 167 | 18 | 148 | 15-Oct-2014 | 1 |
| 41 | 8-Oct-2014 | 15-Oct-2014 | 167 | 17 | 150 | 22-Oct-2014 | 0 |
| 42 | 15-Oct-2014 | 22-Oct-2014 | 167 | 16 | 151 | 29-Oct-2014 | 0 |
| 43 | 22-Oct-2014 | 29-Oct-2014 | 167 | 16 | 150 | 5-Nov-2014 | 1 |
| 44 | 29-Oct-2014 | 5-Nov-2014 | 167 | 17 | 150 | 12-Nov-2014 | 0 |
| 45 | 5-Nov-2014 | 12-Nov-2014 | 167 | 17 | 150 | 19-Nov-2014 | 0 |
| 46 | 12-Nov-2014 | 19-Nov-2014 | 167 | 17 | 149 | 26-Nov-2014 | 1 |
| 47 | 19-Nov-2014 | 26-Nov-2014 | 167 | 18 | 149 | 3-Dec-2014 | 0 |
| 48 | 26-Nov-2014 | 3-Dec-2014 | 167 | 18 | 147 | 10-Dec-2014 | 2 |
| 49 | 3-Dec-2014 | 10-Dec-2014 | 167 | 20 | 146 | 17-Dec-2014 | 1 |
| 50 | 10-Dec-2014 | 17-Dec-2014 | 167 | 21 | 145 | 24-Dec-2014 | 1 |

| | | | | | | | |
|---|---|---|---|---|---|---|---|
| 51 | 17-Dec-2014 | 24-Dec-2014 | 167 | 22 | 145 | 31-Dec-2014 | 0 |
| 52 | 24-Dec-2014 | 31-Dec-2014 | 167 | 22 | 145 | 7-Jan-2015 | 0 |
| 53 | 31-Dec-2014 | 7-Jan-2015 | 167 | 22 | 145 | 14-Jan-2015 | 0 |
| 54 | 7-Jan-2015 | 14-Jan-2015 | 167 | 22 | 145 | 21-Jan-2015 | 0 |
| 55 | 14-Jan-2015 | 21-Jan-2015 | 167 | 22 | 145 | 28-Jan-2015 | 0 |
| 56 | 21-Jan-2015 | 28-Jan-2015 | 167 | 21 | 146 | 4-Feb-2015 | 0 |
| 57 | 28-Jan-2015 | 4-Feb-2015 | 167 | 21 | 146 | 11-Feb-2015 | 0 |
| 58 | 4-Feb-2015 | 11-Feb-2015 | 167 | 21 | 145 | 18-Feb-2015 | 1 |
| 59 | 11-Feb-2015 | 18-Feb-2015 | 167 | 22 | 145 | 25-Feb-2015 | 0 |
| 60 | 18-Feb-2015 | 25-Feb-2015 | 167 | 21 | 146 | 4-Mar-2015 | 0 |
| 61 | 25-Feb-2015 | 4-Mar-2015 | 167 | 21 | 146 | 11-Mar-2015 | 0 |
| 62 | 4-Mar-2015 | 11-Mar-2015 | 167 | 20 | 146 | 18-Mar-2015 | 1 |
| 63 | 11-Mar-2015 | 18-Mar-2015 | 167 | 21 | 146 | 25-Mar-2015 | 0 |
| 64 | 18-Mar-2015 | 25-Mar-2015 | 167 | 21 | 144 | 1-Apr-2015 | 2 |
| 65 | 25-Mar-2015 | 1-Apr-2015 | 167 | 23 | 144 | 8-Apr-2015 | 0 |
| 66 | 1-Apr-2015 | 8-Apr-2015 | 167 | 23 | 140 | 15-Apr-2015 | 4 |
| 67 | 8-Apr-2015 | 15-Apr-2015 | 167 | 27 | 140 | 22-Apr-2015 | 0 |
| 68 | 15-Apr-2015 | 22-Apr-2015 | 167 | 27 | 139 | 29-Apr-2015 | 1 |
| 69 | 22-Apr-2015 | 29-Apr-2015 | 167 | 28 | 138 | 6-May-2015 | 1 |
| 70 | 29-Apr-2015 | 6-May-2015 | 167 | 27 | 140 | 13-May-2015 | 0 |
| 71 | 6-May-2015 | 13-May-2015 | 167 | 27 | 140 | 20-May-2015 | 0 |
| 72 | 13-May-2015 | 20-May-2015 | 167 | 27 | 139 | 27-May-2015 | 1 |
| 73 | 20-May-2015 | 27-May-2015 | 167 | 28 | 139 | 3-Jun-2015 | 0 |
| 74 | 27-May-2015 | 3-Jun-2015 | 167 | 28 | 138 | 10-Jun-2015 | 1 |
| 75 | 3-Jun-2015 | 10-Jun-2015 | 167 | 28 | 139 | 17-Jun-2015 | 0 |
| 76 | 10-Jun-2015 | 17-Jun-2015 | 167 | 28 | 139 | 24-Jun-2015 | 0 |
| 77 | 17-Jun-2015 | 24-Jun-2015 | 167 | 28 | 138 | 1-Jul-2015 | 1 |
| 78 | 24-Jun-2015 | 1-Jul-2015 | 167 | 28 | 133 | 8-Jul-2015 | 6 |
| 79 | 1-Jul-2015 | 8-Jul-2015 | 167 | 33 | 133 | 15-Jul-2015 | 1 |
| 80 | 8-Jul-2015 | 15-Jul-2015 | 167 | 31 | 135 | 22-Jul-2015 | 1 |
| 81 | 15-Jul-2015 | 22-Jul-2015 | 167 | 32 | 135 | 29-Jul-2015 | 0 |
| 82 | 22-Jul-2015 | 29-Jul-2015 | 167 | 30 | 137 | 5-Aug-2015 | 0 |
| 83 | 29-Jul-2015 | 5-Aug-2015 | 167 | 30 | 135 | 12-Aug-2015 | 2 |
| 84 | 5-Aug-2015 | 12-Aug-2015 | 167 | 32 | 135 | 19-Aug-2015 | 0 |
| 85 | 12-Aug-2015 | 19-Aug-2015 | 167 | 32 | 135 | 26-Aug-2015 | 0 |
| 86 | 19-Aug-2015 | 26-Aug-2015 | 167 | 32 | 135 | 2-Sep-2015 | 0 |
| 87 | 26-Aug-2015 | 2-Sep-2015 | 167 | 32 | 135 | 9-Sep-2015 | 0 |
| 88 | 2-Sep-2015 | 9-Sep-2015 | 167 | 32 | 134 | 16-Sep-2015 | 1 |
| 89 | 9-Sep-2015 | 16-Sep-2015 | 167 | 33 | 134 | 23-Sep-2015 | 0 |
| 90 | 16-Sep-2015 | 23-Sep-2015 | 167 | 33 | 134 | 30-Sep-2015 | 0 |
| 91 | 23-Sep-2015 | 30-Sep-2015 | 167 | 33 | 134 | 7-Oct-2015 | 0 |
| 92 | 30-Sep-2015 | 7-Oct-2015 | 167 | 32 | 135 | 14-Oct-2015 | 0 |
| 93 | 7-Oct-2015 | 14-Oct-2015 | 167 | 32 | 135 | 21-Oct-2015 | 0 |
| 94 | 14-Oct-2015 | 21-Oct-2015 | 167 | 27 | 139 | 28-Oct-2015 | 1 |
| 95 | 21-Oct-2015 | 28-Oct-2015 | 167 | 28 | 137 | 4-Nov-2015 | 2 |
| 96 | 28-Oct-2015 | 4-Nov-2015 | 167 | 30 | 136 | 11-Nov-2015 | 1 |
| 97 | 4-Nov-2015 | 11-Nov-2015 | 167 | 31 | 133 | 18-Nov-2015 | 3 |
| 98 | 11-Nov-2015 | 18-Nov-2015 | 167 | 34 | 132 | 25-Nov-2015 | 1 |
| 99 | 18-Nov-2015 | 25-Nov-2015 | 167 | 35 | 131 | 2-Dec-2015 | 1 |
| 100 | 25-Nov-2015 | 2-Dec-2015 | 167 | 36 | 128 | 9-Dec-2015 | 3 |
| 101 | 2-Dec-2015 | 9-Dec-2015 | 167 | 37 | 129 | 16-Dec-2015 | 1 |
| 102 | 9-Dec-2015 | 16-Dec-2015 | 167 | 38 | 128 | 23-Dec-2015 | 1 |
| 103 | 16-Dec-2015 | 23-Dec-2015 | 167 | 39 | 128 | 30-Dec-2015 | 0 |
| 104 | 23-Dec-2015 | 30-Dec-2015 | 167 | 39 | 126 | 6-Jan-2016 | 2 |
| 105 | 30-Dec-2015 | 6-Jan-2016 | 167 | 40 | 126 | 13-Jan-2016 | 1 |
| 106 | 6-Jan-2016 | 13-Jan-2016 | 167 | 41 | 125 | 20-Jan-2016 | 1 |
| 107 | 13-Jan-2016 | 20-Jan-2016 | 167 | 42 | 124 | 27-Jan-2016 | 1 |
| 108 | 20-Jan-2016 | 27-Jan-2016 | 167 | 43 | 124 | 3-Feb-2016 | 0 |
| 109 | 27-Jan-2016 | 3-Feb-2016 | 167 | 43 | 124 | 10-Feb-2016 | 0 |
| 110 | 3-Feb-2016 | 10-Feb-2016 | 167 | 43 | 124 | 17-Feb-2016 | 0 |

| | | | | | | | |
|---|---|---|---|---|---|---|---|
| 111 | 10-Feb-2016 | 17-Feb-2016 | 167 | 42 | 124 | 24-Feb-2016 | 1 |
| 112 | 17-Feb-2016 | 24-Feb-2016 | 167 | 42 | 125 | 2-Mar-2016 | 0 |
| 113 | 24-Feb-2016 | 2-Mar-2016 | 167 | 42 | 123 | 9-Mar-2016 | 2 |
| 114 | 2-Mar-2016 | 9-Mar-2016 | 167 | 44 | 121 | 16-Mar-2016 | 2 |
| 115 | 9-Mar-2016 | 16-Mar-2016 | 167 | 46 | 121 | 23-Mar-2016 | 0 |
| 116 | 16-Mar-2016 | 23-Mar-2016 | 167 | 46 | 121 | 30-Mar-2016 | 0 |
| 117 | 23-Mar-2016 | 30-Mar-2016 | 167 | 46 | 120 | 6-Apr-2016 | 1 |
| 118 | 30-Mar-2016 | 6-Apr-2016 | 167 | 46 | 120 | 13-Apr-2016 | 1 |
| 119 | 6-Apr-2016 | 13-Apr-2016 | 167 | 42 | 124 | 20-Apr-2016 | 1 |
| 120 | 13-Apr-2016 | 20-Apr-2016 | 167 | 43 | 124 | 27-Apr-2016 | 0 |
| 121 | 20-Apr-2016 | 27-Apr-2016 | 167 | 43 | 124 | 4-May-2016 | 0 |
| 122 | 27-Apr-2016 | 4-May-2016 | 167 | 42 | 125 | 11-May-2016 | 0 |
| 123 | 4-May-2016 | 11-May-2016 | 167 | 40 | 126 | 18-May-2016 | 1 |
| 124 | 11-May-2016 | 18-May-2016 | 167 | 39 | 128 | 25-May-2016 | 0 |
| 125 | 18-May-2016 | 25-May-2016 | 167 | 39 | 127 | 1-Jun-2016 | 1 |
| 126 | 25-May-2016 | 1-Jun-2016 | 167 | 39 | 127 | 8-Jun-2016 | 1 |
| 127 | 1-Jun-2016 | 8-Jun-2016 | 167 | 39 | 128 | 15-Jun-2016 | 0 |
| 128 | 8-Jun-2016 | 15-Jun-2016 | 167 | 38 | 128 | 22-Jun-2016 | 1 |
| 129 | 15-Jun-2016 | 22-Jun-2016 | 167 | 39 | 128 | 29-Jun-2016 | 0 |
| 130 | 22-Jun-2016 | 29-Jun-2016 | 167 | 37 | 129 | 6-Jul-2016 | 1 |
| 131 | 29-Jun-2016 | 6-Jul-2016 | 167 | 37 | 129 | 13-Jul-2016 | 1 |
| 132 | 6-Jul-2016 | 13-Jul-2016 | 167 | 38 | 129 | 20-Jul-2016 | 0 |
| 133 | 13-Jul-2016 | 20-Jul-2016 | 167 | 38 | 129 | 27-Jul-2016 | 0 |
| 134 | 20-Jul-2016 | 27-Jul-2016 | 167 | 35 | 131 | 3-Aug-2016 | 1 |
| 135 | 27-Jul-2016 | 3-Aug-2016 | 167 | 35 | 132 | 10-Aug-2016 | 0 |
| 136 | 3-Aug-2016 | 10-Aug-2016 | 167 | 33 | 134 | 17-Aug-2016 | 0 |
| 137 | 10-Aug-2016 | 17-Aug-2016 | 167 | 32 | 135 | 24-Aug-2016 | 0 |
| 138 | 17-Aug-2016 | 24-Aug-2016 | 167 | 30 | 137 | 31-Aug-2016 | 0 |
| 139 | 24-Aug-2016 | 31-Aug-2016 | 167 | 28 | 139 | 7-Sep-2016 | 0 |
| 140 | 31-Aug-2016 | 7-Sep-2016 | 167 | 28 | 138 | 14-Sep-2016 | 1 |
| 141 | 7-Sep-2016 | 14-Sep-2016 | 167 | 29 | 138 | 21-Sep-2016 | 0 |
| 142 | 14-Sep-2016 | 21-Sep-2016 | 167 | 29 | 136 | 28-Sep-2016 | 2 |
| 143 | 21-Sep-2016 | 28-Sep-2016 | 167 | 30 | 137 | 5-Oct-2016 | 0 |
| 144 | 28-Sep-2016 | 5-Oct-2016 | 167 | 30 | 137 | 12-Oct-2016 | 0 |
| 145 | 5-Oct-2016 | 12-Oct-2016 | 167 | 30 | 136 | 19-Oct-2016 | 1 |
| 146 | 12-Oct-2016 | 19-Oct-2016 | 167 | 31 | 136 | 26-Oct-2016 | 0 |
| 147 | 19-Oct-2016 | 26-Oct-2016 | 167 | 29 | 137 | 2-Nov-2016 | 1 |
| 148 | 26-Oct-2016 | 2-Nov-2016 | 167 | 30 | 136 | 9-Nov-2016 | 1 |
| 149 | 2-Nov-2016 | 9-Nov-2016 | 167 | 31 | 136 | 16-Nov-2016 | 0 |
| 150 | 9-Nov-2016 | 16-Nov-2016 | 167 | 31 | 136 | 23-Nov-2016 | 0 |
| 151 | 16-Nov-2016 | 23-Nov-2016 | 167 | 31 | 135 | 30-Nov-2016 | 1 |
| 152 | 23-Nov-2016 | 30-Nov-2016 | 167 | 32 | 133 | 7-Dec-2016 | 2 |
| 153 | 30-Nov-2016 | 7-Dec-2016 | 167 | 33 | 132 | 14-Dec-2016 | 2 |
| 154 | 7-Dec-2016 | 14-Dec-2016 | 167 | 27 | 140 | 21-Dec-2016 | 0 |
| 155 | 14-Dec-2016 | 21-Dec-2016 | 167 | 27 | 139 | 28-Dec-2016 | 1 |
| 156 | 21-Dec-2016 | 28-Dec-2016 | 167 | 28 | 138 | 4-Jan-2017 | 1 |
| 157 | 28-Dec-2016 | 4-Jan-2017 | 167 | 29 | 138 | 11-Jan-2017 | 0 |
| 158 | 4-Jan-2017 | 11-Jan-2017 | 167 | 29 | 138 | 18-Jan-2017 | 0 |
| 159 | 11-Jan-2017 | 18-Jan-2017 | 167 | 29 | 137 | 25-Jan-2017 | 1 |
| 160 | 18-Jan-2017 | 25-Jan-2017 | 167 | 30 | 137 | 1-Feb-2017 | 0 |
| 161 | 25-Jan-2017 | 1-Feb-2017 | 167 | 30 | 136 | 8-Feb-2017 | 1 |
| 162 | 1-Feb-2017 | 8-Feb-2017 | 167 | 31 | 136 | 15-Feb-2017 | 0 |
| 163 | 8-Feb-2017 | 15-Feb-2017 | 167 | 31 | 135 | 22-Feb-2017 | 1 |
| 164 | 15-Feb-2017 | 22-Feb-2017 | 167 | 32 | 134 | 1-Mar-2017 | 1 |
| 165 | 22-Feb-2017 | 1-Mar-2017 | 167 | 32 | 134 | 8-Mar-2017 | 1 |
| 166 | 1-Mar-2017 | 8-Mar-2017 | 167 | 32 | 135 | 15-Mar-2017 | 0 |
| 167 | 8-Mar-2017 | 15-Mar-2017 | 167 | 31 | 136 | 22-Mar-2017 | 0 |
| 168 | 15-Mar-2017 | 22-Mar-2017 | 167 | 31 | 135 | 29-Mar-2017 | 1 |
| 169 | 22-Mar-2017 | 29-Mar-2017 | 167 | 32 | 134 | 5-Apr-2017 | 1 |
| 170 | 29-Mar-2017 | 5-Apr-2017 | 167 | 32 | 129 | 12-Apr-2017 | 6 |

| | | | | | | | |
|---|---|---|---|---|---|---|---|
| 171 | 5-Apr-2017 | 12-Apr-2017 | 167 | 37 | 129 | 19-Apr-2017 | 1 |
| 172 | 12-Apr-2017 | 19-Apr-2017 | 167 | 38 | 128 | 26-Apr-2017 | 1 |
| 173 | 19-Apr-2017 | 26-Apr-2017 | 167 | 39 | 127 | 3-May-2017 | 1 |
| 174 | 26-Apr-2017 | 3-May-2017 | 167 | 39 | 128 | 10-May-2017 | 0 |
| 175 | 3-May-2017 | 10-May-2017 | 167 | 38 | 129 | 17-May-2017 | 0 |
| 176 | 10-May-2017 | 17-May-2017 | 167 | 37 | 129 | 24-May-2017 | 1 |
| 177 | 17-May-2017 | 24-May-2017 | 167 | 36 | 131 | 31-May-2017 | 0 |
| 178 | 24-May-2017 | 31-May-2017 | 167 | 36 | 131 | 7-Jun-2017 | 0 |
| 179 | 31-May-2017 | 7-Jun-2017 | 167 | 35 | 132 | 14-Jun-2017 | 0 |
| 180 | 7-Jun-2017 | 14-Jun-2017 | 167 | 35 | 130 | 21-Jun-2017 | 2 |
| 181 | 14-Jun-2017 | 21-Jun-2017 | 167 | 36 | 131 | 28-Jun-2017 | 0 |
| 182 | 21-Jun-2017 | 28-Jun-2017 | 167 | 36 | 129 | 5-Jul-2017 | 2 |
| 183 | 28-Jun-2017 | 5-Jul-2017 | 167 | 38 | 128 | 12-Jul-2017 | 1 |
| 184 | 5-Jul-2017 | 12-Jul-2017 | 167 | 39 | 128 | 19-Jul-2017 | 0 |
| 185 | 12-Jul-2017 | 19-Jul-2017 | 167 | 39 | 126 | 26-Jul-2017 | 2 |
| 186 | 19-Jul-2017 | 26-Jul-2017 | 167 | 41 | 123 | 2-Aug-2017 | 3 |
| 187 | 26-Jul-2017 | 2-Aug-2017 | 167 | 42 | 125 | 9-Aug-2017 | 0 |
| 188 | 2-Aug-2017 | 9-Aug-2017 | 167 | 38 | 129 | 16-Aug-2017 | 0 |
| 189 | 9-Aug-2017 | 16-Aug-2017 | 167 | 37 | 130 | 23-Aug-2017 | 0 |
| 190 | 16-Aug-2017 | 23-Aug-2017 | 167 | 36 | 131 | 30-Aug-2017 | 0 |
| 191 | 23-Aug-2017 | 30-Aug-2017 | 167 | 35 | 132 | 6-Sep-2017 | 0 |
| 192 | 30-Aug-2017 | 6-Sep-2017 | 167 | 35 | 132 | 13-Sep-2017 | 0 |
| 193 | 6-Sep-2017 | 13-Sep-2017 | 167 | 35 | 131 | 20-Sep-2017 | 1 |
| 194 | 13-Sep-2017 | 20-Sep-2017 | 167 | 35 | 131 | 27-Sep-2017 | 1 |
| 195 | 20-Sep-2017 | 27-Sep-2017 | 167 | 36 | 130 | 4-Oct-2017 | 1 |
| 196 | 27-Sep-2017 | 4-Oct-2017 | 167 | 37 | 129 | 11-Oct-2017 | 1 |
| 197 | 4-Oct-2017 | 11-Oct-2017 | 167 | 37 | 129 | 18-Oct-2017 | 1 |
| 198 | 11-Oct-2017 | 18-Oct-2017 | 167 | 36 | 130 | 25-Oct-2017 | 1 |
| 199 | 18-Oct-2017 | 25-Oct-2017 | 167 | 37 | 129 | 1-Nov-2017 | 1 |
| 200 | 25-Oct-2017 | 1-Nov-2017 | 167 | 38 | 127 | 8-Nov-2017 | 2 |
| 201 | 1-Nov-2017 | 8-Nov-2017 | 167 | 40 | 126 | 15-Nov-2017 | 1 |
| 202 | 8-Nov-2017 | 15-Nov-2017 | 167 | 41 | 125 | 22-Nov-2017 | 1 |
| 203 | 15-Nov-2017 | 22-Nov-2017 | 167 | 42 | 123 | 29-Nov-2017 | 2 |
| 204 | 22-Nov-2017 | 29-Nov-2017 | 167 | 42 | 124 | 6-Dec-2017 | 1 |
| 205 | 29-Nov-2017 | 6-Dec-2017 | 167 | 41 | 125 | 13-Dec-2017 | 1 |
| 206 | 6-Dec-2017 | 13-Dec-2017 | 167 | 42 | 122 | 20-Dec-2017 | 3 |
| 207 | 13-Dec-2017 | 20-Dec-2017 | 167 | 45 | 121 | 27-Dec-2017 | 1 |
| 208 | 20-Dec-2017 | 27-Dec-2017 | 167 | 46 | 117 | 3-Jan-2018 | 4 |
| 209 | 27-Dec-2017 | 3-Jan-2018 | 167 | 50 | 117 | 10-Jan-2018 | 0 |
| 210 | 3-Jan-2018 | 10-Jan-2018 | 167 | 48 | 117 | 17-Jan-2018 | 2 |
| 211 | 10-Jan-2018 | 17-Jan-2018 | 167 | 50 | 115 | 24-Jan-2018 | 2 |
| 212 | 17-Jan-2018 | 24-Jan-2018 | 167 | 52 | 113 | 31-Jan-2018 | 2 |
| 213 | 24-Jan-2018 | 31-Jan-2018 | 167 | 54 | 112 | 7-Feb-2018 | 1 |
| 214 | 31-Jan-2018 | 7-Feb-2018 | 167 | 54 | 113 | 14-Feb-2018 | 0 |
| 215 | 7-Feb-2018 | 14-Feb-2018 | 167 | 54 | 113 | 21-Feb-2018 | 0 |
| 216 | 14-Feb-2018 | 21-Feb-2018 | 167 | 54 | 112 | 28-Feb-2018 | 1 |
| 217 | 21-Feb-2018 | 28-Feb-2018 | 167 | 55 | 112 | 7-Mar-2018 | 0 |
| 218 | 28-Feb-2018 | 7-Mar-2018 | 167 | 55 | 111 | 14-Mar-2018 | 1 |
| 219 | 7-Mar-2018 | 14-Mar-2018 | 167 | 55 | 112 | 21-Mar-2018 | 0 |
| 220 | 14-Mar-2018 | 21-Mar-2018 | 167 | 55 | 112 | 28-Mar-2018 | 0 |
| 221 | 21-Mar-2018 | 28-Mar-2018 | 167 | 54 | 111 | 4-Apr-2018 | 2 |
| 222 | 28-Mar-2018 | 4-Apr-2018 | 167 | 56 | 111 | 11-Apr-2018 | 0 |
| 223 | 4-Apr-2018 | 11-Apr-2018 | 167 | 56 | 109 | 18-Apr-2018 | 2 |
| 224 | 11-Apr-2018 | 18-Apr-2018 | 167 | 58 | 109 | 25-Apr-2018 | 0 |
| 225 | 18-Apr-2018 | 25-Apr-2018 | 167 | 58 | 109 | 2-May-2018 | 0 |
| 226 | 25-Apr-2018 | 2-May-2018 | 167 | 58 | 109 | 9-May-2018 | 0 |
| 227 | 2-May-2018 | 9-May-2018 | 167 | 58 | 109 | 16-May-2018 | 0 |
| 228 | 9-May-2018 | 16-May-2018 | 167 | 57 | 110 | 23-May-2018 | 0 |
| 229 | 16-May-2018 | 23-May-2018 | 167 | 54 | 113 | 30-May-2018 | 0 |
| 230 | 23-May-2018 | 30-May-2018 | 167 | 54 | 112 | 6-Jun-2018 | 1 |

| | | | | | | | |
|---|---|---|---|---|---|---|---|
| 231 | 30-May-2018 | 6-Jun-2018 | 167 | 53 | 113 | 13-Jun-2018 | 1 |
| 232 | 6-Jun-2018 | 13-Jun-2018 | 167 | 54 | 111 | 20-Jun-2018 | 2 |
| 233 | 13-Jun-2018 | 20-Jun-2018 | 167 | 56 | 110 | 27-Jun-2018 | 1 |
| 234 | 20-Jun-2018 | 27-Jun-2018 | 167 | 57 | 110 | 4-Jul-2018 | 0 |
| 235 | 27-Jun-2018 | 4-Jul-2018 | 167 | 57 | 110 | 11-Jul-2018 | 0 |
| 236 | 4-Jul-2018 | 11-Jul-2018 | 167 | 57 | 110 | 18-Jul-2018 | 0 |
| 237 | 11-Jul-2018 | 18-Jul-2018 | 167 | 56 | 111 | 25-Jul-2018 | 0 |
| 238 | 18-Jul-2018 | 25-Jul-2018 | 167 | 56 | 110 | 1-Aug-2018 | 1 |
| 239 | 25-Jul-2018 | 1-Aug-2018 | 167 | 52 | 115 | 8-Aug-2018 | 0 |
| 240 | 1-Aug-2018 | 8-Aug-2018 | 167 | 52 | 115 | 15-Aug-2018 | 0 |
| 241 | 8-Aug-2018 | 15-Aug-2018 | 167 | 51 | 116 | 22-Aug-2018 | 0 |
| 242 | 15-Aug-2018 | 22-Aug-2018 | 167 | 48 | 118 | 29-Aug-2018 | 1 |
| 243 | 22-Aug-2018 | 29-Aug-2018 | 167 | 45 | 122 | 5-Sep-2018 | 0 |
| 244 | 29-Aug-2018 | 5-Sep-2018 | 167 | 45 | 122 | 12-Sep-2018 | 0 |
| 245 | 5-Sep-2018 | 12-Sep-2018 | 167 | 45 | 122 | 19-Sep-2018 | 0 |
| 246 | 12-Sep-2018 | 19-Sep-2018 | 167 | 42 | 125 | 26-Sep-2018 | 0 |
| 247 | 19-Sep-2018 | 26-Sep-2018 | 167 | 41 | 126 | 3-Oct-2018 | 0 |
| 248 | 26-Sep-2018 | 3-Oct-2018 | 167 | 37 | 130 | 10-Oct-2018 | 0 |
| 249 | 3-Oct-2018 | 10-Oct-2018 | 167 | 36 | 131 | 17-Oct-2018 | 0 |
| 250 | 10-Oct-2018 | 17-Oct-2018 | 167 | 36 | 131 | 24-Oct-2018 | 0 |
| 251 | 17-Oct-2018 | 24-Oct-2018 | 167 | 36 | 129 | 31-Oct-2018 | 2 |
| 252 | 24-Oct-2018 | 31-Oct-2018 | 167 | 36 | 130 | 7-Nov-2018 | 1 |
| 253 | 31-Oct-2018 | 7-Nov-2018 | 167 | 36 | 131 | 14-Nov-2018 | 0 |
| 254 | 7-Nov-2018 | 14-Nov-2018 | 167 | 35 | 131 | 21-Nov-2018 | 1 |
| 255 | 14-Nov-2018 | 21-Nov-2018 | 167 | 36 | 129 | 28-Nov-2018 | 2 |
| 256 | 21-Nov-2018 | 28-Nov-2018 | 167 | 36 | 129 | 5-Dec-2018 | 2 |
| 257 | 28-Nov-2018 | 5-Dec-2018 | 167 | 38 | 129 | 12-Dec-2018 | 0 |
| 258 | 5-Dec-2018 | 12-Dec-2018 | 167 | 38 | 129 | 19-Dec-2018 | 0 |
| 259 | 12-Dec-2018 | 19-Dec-2018 | 167 | 38 | 129 | 26-Dec-2018 | 0 |

* Number of participant farms could vary by week. The risk was estimated for the participant farms in the Morrison Swine Health Monitoring Project (MSHMP) of the University of Minnesota (https://vetmed.umn.edu/centers-programs/swine-program/outreach-leman-mshmp/mshmp, accessed on 20 March 2020).
